# Predicting individual shelter dog behaviour after adoption using longitudinal behavioural assessment: a hierarchical Bayesian approach

**DOI:** 10.1101/2021.05.13.443965

**Authors:** Conor Goold, Ruth C. Newberry

## Abstract

Predicting the behaviour of shelter dogs after adoption is an important, but difficult, endeavour. Differences between shelter and post-adoption environments, between- and within-individual heterogeneity in behaviour, uncertainty in behavioural predictions and measurement error all hinder the accurate assessment of future behaviour. This study integrates 1) a longitudinal behavioural assessment with 2) a novel joint hierarchical Bayesian mixture model that accounts for individual variation, missing data and measurement error to predict behaviour post-adoption. We analysed shelter observations (> 28,000 records) and post-adoption reports (from telephone surveys) on the behaviour of 241 dogs across eight contexts. Dog behaviour at the shelter correlated positively with behaviour post-adoption within contexts (*r* = 0.38; 95% highest density interval: [0.20, 0.55]), and behavioural repeatability was approximately 20% higher post-adoption than at the shelter for behaviour within contexts. Although measurement error was higher post-adoption than at the shelter, we found few differences in individual-level, latent probabilities of different behaviours post-adoption versus at the shelter. This good predictive ability was aided by accurate representation of uncertainty in individual-level predictions. We conclude that longitudinal assessment paired with a sufficient inferential framework to model latent behavioural profiles with uncertainty enables reasonably accurate estimation of post-adoption behaviour.

## 1 Introduction

Prediction is a defining, yet challenging, goal of science (Hofstadter 1951), uniting a diverse range of disciplines including climate research, personality psychology, social policy, medicine, conservation biology, and complex systems science (Sarewitz and Pielke 1999). More specifically, research aimed at predicting the future behaviour of domestic animals is of both basic interest in evaluating our understanding of biological processes and applied interest for decision-making relating to animal use and human safety. One important application concerns accurately predicting the future behaviour of domestic dogs (*Canis familiaris*), which represents a challenge for professional organisations, dog breeders, and dog owners alike. In particular, animal shelters have the difficult task of assessing how dogs will behave in a variety of circumstances from limited behavioural information and making decisions about the suitability of those dogs for placement into new homes.

Animal shelters frequently employ standardised behaviour tests that evaluate specific behavioural or *personality* traits through reconstructions of situations relevant to life outside the shelter environment (van der Borg et al. 1991; Marston and Bennett 2003; Mornement et al. 2010; Taylor and Mills 2006; Rayment et al. 2015; Clay et al. 2020a). For example, to assess a dog’s level of aggressiveness around food (i.e. food guarding), shelter staff may present a dog with a bowl of food and record the dog’s response to a plastic hand approaching, touching or trying to remove the food bowl (Mohan-Gibbons et al. 2012; Mohan-Gibbons et al. 2018; Marder et al. 2013). Such tests are usually conducted once, soon after arrival at the shelter, yielding an overall score used to help determine a dog’s suitability for adoption. However, criticisms have been raised about the ability of these protocols to provide accurate predictions of future dog behaviour. For example, the feasibility of carrying out a battery of such behaviour tests with high reliability and standardisation in the time-constrained shelter environment is limited given that test batteries often require at least an hour per dog (van der Borg et al. 1991; Mornement et al. 2009; although see Poulsen et al. (2010) for a shorter protocol). Moreover, a high number of false positives have been recorded for food guarding behaviour (i.e. displaying food aggression during shelter evaluations but not in the new home; see Mohan-Gibbons et al. 2012; Marder et al. 2013), leading to calls to abandon standardised food tests given that dog return rates to shelters do not appear to increase if the tests are not conducted (Mohan-Gibbons et al. 2018). Behaviour in other contexts might be similarly difficult to predict post-adoption (Christensen et al. 2007; Mornement et al. 2015; although see Bollen and Horowitz (2008)).

Ways to improve predictions about individual dog behaviour post-adoption are not entirely clear. While emphasis has been placed on improving the reliability and validity of shelter dog tests, progress is hampered by a lack of clarity of what those terms mean (Patronek et al. 2019; Rayment et al. 2015), which is also a problem for the wider scientific community (e.g. see Borsboom et al. 2004; Borsboom et al. 2009; Maul et al. 2016). For instance, Patronek and Bradley (2016) demonstrate that even with high sensitivity and specificity, the probability of a dog showing aggression in the new home after a positive test in the shelter is unlikely to be higher than 50% and is probably closer to ∼ 30% (see also Patronek et al. 2019). Tests reported to predict post-adoption behaviour successfully (e.g. Valsecchi et al. 2011; Poulsen et al. 2010; Bollen and Horowitz 2008) have been supported based on the statistical significance of linear associations (e.g. significant correlations or regression coefficients) between shelter and post-adoption behaviour (i.e. predictive ‘validity’; Patronek et al. 2019). In contrast, arguments against the efficacy of behavioural tests highlight evidence of low predictive ‘ability’ for individual dogs (i.e. estimates of true/false positive and negative rates; Patronek et al. 2019).

As an alternative to standardised testing, several authors and organisations have emphasised the collection of daily behavioural observation records from dogs in shelters, in addition to other information (e.g. foster reports, pre-surrender interviews). A holistic profile of each dog’s behaviour and welfare needs is then formulated and used to inform adoption decisions (Patronek et al. 2019; ASPCA 2018; Rayment et al. 2015; Mornement et al. 2015; Clay et al. 2020b). Such longitudinal approaches refrain from crudely labelling individual dogs as, for example, aggressive or non-aggressive based on a single behavioural testing outcome. However, research is still scarce on how best to implement this approach, which requires summarising swathes of information generated on each dog, in a manner that is of practical use for shelters.

We have previously reported on the behaviour of dogs while in shelters based on data collected using the longitudinal assessment methodology implemented by a large UK shelter organisation (Goold and Newberry 2017b; Goold and Newberry 2017a). The assessment relies on the spontaneous reporting of behavioural observations made by shelter staff during everyday contexts (e.g. walking past the dog’s kennel, putting a food bowl into the kennel, clipping on the lead or touching the collar). From these accumulated reports, Goold and Newberry (2017a) used the framework of behavioural reaction norms (Dingemanse et al. 2010; Cleasby et al. 2015) to partition individual variation in the behaviour of over 3,000 dogs when around unknown people (from almost 20,000 behavioural reports in total) into three components: personality (i.e. difference in average behaviour between individuals), plasticity (i.e. within-individual behavioural change over time) and predictability (i.e. within-individual residual variance). Accounting for all three components improved the estimated out-of-sample predictive accuracy of the statistical models.

While accounting for individual variation in behaviour across time at the shelter is important, making predictions about dog behaviour post-adoption based on longitudinal information about behaviour at the shelter brings its own challenges. Deciding on what statistical moments (e.g. location, scale) of the longitudinal shelter data should predict post-adoption behaviour is not obvious. Diagnostic statistics such as false positives or false negatives lose clarity because longitudinal assessments acknowledge that dog behaviour can change depending on time and context (Goold and Newberry 2017a). Inferential tools that can estimate the individual-level, latent probabilities of dogs showing different behaviours at different time points are required, rather than focusing on single occurrences of behaviours. The reality of collecting observational data in busy shelter environments also means that data may present significant patterns of non-ignorable missingness and substantially unbalanced repeated measures, and be subject to more measurement error than standardised assessments. Altogether, determining the predictive value of ‘on-going’ shelter assessments requires careful analysis of longitudinal data using statistical models that appropriately account for the data-generating processes.

The goal of the present study was to assess the correspondence between behaviour shown at the shelter, collected using the longitudinal assessment methodology, and the behaviour reported during telephone interviews post-adoption. Our secondary goal was to apply a novel application of joint Bayesian hierarchical mixture modelling to compare predictions about dog behaviour in shelters and post-adoption. Importantly, our approach accounts for i) individual variation in both average behaviour (i.e. personality) and behavioural change (i.e. plasticity; due to only two post-adoption interviews, predictability was not estimated) across shelter and post-adoption time periods, ii) missing data and their potential correlates, and iii) measurement error in the behavioural reports. We report on predictive validity by estimating correlations between personality and plasticity between shelter and post-adoption time periods. Predictive ability is examined by comparing model predictions for each dog’s average behaviour in the shelter and post-adoption to determine whether dogs exhibited credible changes in their behaviour. We present specific examples of individual-level behaviour in our results and discussion.

## 2 Materials & Methods

### 2.1 Subjects

Behavioural data were gathered retrospectively from new owners of 265 dogs adopted from Battersea Dogs and Cats Home, United Kingdom between May 2016 and May 2017. Details of the shelter environment can be found in Goold and Newberry (2017b) and Goold and Newberry (2017a). Of the 265 dogs, only 241 dogs could be matched with behavioural and demographic data from the shelter’s database. Demographic information about these dogs is presented in Table 1. These dogs ranged from approximately 8 weeks old to 13 years old (estimated by shelter staff when unknown) and were generally of unknown heritage (i.e. not in a breed registry). They were rehomed from different shelter facilities. Demographic information on the new owners was collected but not analysed here due to inconsistencies in the responses. The original dog identification numbers assigned to dogs by the shelter were removed prior to analysis.

**Table 1:** *Dog demographic variables collected at the shelter*.

| Demographic variable | Mean (SD) or $n$ dogs |
| --- | --- |
| Number of observations per dog | 140.7 (224.3) |
| Days at the shelter per dog | 32.1 (41.4) |
| Estimated age at departure from shelter (years) | 3.6 (2.8) |
| Weight (kg; last record at shelter) | 16.5 (9.6) |
| Source (before surveys): gift/stray/return ($n$) | 164/51/26 |
| Sex: female/male ($n$) | 102/139 |
| Neutered: before arrival/at shelter/not/missing ($n$) | 84/145/11/1 |
| Rehoming centre: London/Old Windsor/Brands Hatch ($n$) | 71/93/77 |

### 2.2 Data collection

#### 2.2.1 Shelter behaviour

As previously described by Goold and Newberry (2017a) and Goold and Newberry (2017b), shelter employees observed dogs in a variety of naturally occurring contexts and recorded each dog’s behaviour using a context-specific list of mutually exclusive behavioural codes. The analysis presented here focussed on eight core ‘on-site’ contexts (Table 2). The *Interactions with dogs* context was a combination of the original *Interactions with female* and *male dogs* contexts, respectively, because the post-adoption data did not distinguish between interactions with male and female dogs (as adopters may not have been fully aware of the sex of other dogs that their dog met). Observation records were entered as often as possible given the frequency at which the context occurred and time constraints on shelter staff entering data. We considered any day that a dog did not receive an observation record in a context as a missing observation so that patterns of missingness could be modelled (see Table 2 for frequency of missing daily observation records). In total, there were 28, 445 complete observation records and 33, 507 missing observation records for the 241 studied dogs in the shelter dataset.

There were between ten and sixteen possible behavioural codes depending on the context, arranged on a scale of perceived ease of adoption or desirability to adopters (see Supplementary Materials for the full list of behavioural codes). The longitudinal assessment further categorised the behavioural codes into green, amber and red categories: green codes indicated that the dog’s behaviour was suitable for immediate adoption and/or would be easy for all owners to manage once adopted (e.g. the dog was friendly or excited when meeting new people), amber codes indicated that some training or management might be needed to manage or improve behaviour (e.g. the dog was nervous when meeting new people and slow to meet with them), and red codes that the dog needed an individualised training program and improvement in behaviour to facilitate adoption or would be the most difficult for adopters to manage (e.g. the dog showed aggression when meeting unfamiliar people). A dog’s suitability for adoption was decided based on behaviour across all contexts and days after arrival.

The behavioural scores were analysed using the green, amber and red ordinal scale, rather than using the individual behavioural codes of each ethogram. This was chosen: i) to place the behaviour records across contexts on a comparable scale, allowing all the data to be analysed within the same statistical model; ii) to capture the main variation in the data given that some individual codes were seldom used; iii) because previous analysis indicated higher consensus among shelter staff when rating behaviour using the green, amber and red codes than the individual codes (see Goold and Newberry 2017a); and iv) it was more practically relevant because the shelter was interested to find out if their assessments at the shelter differed greatly from a dog’s behaviour post-adoption (i.e. there was a change from green to red codes), but not necessarily if there was a change between codes within the same colour category. When more than one record was made of the same dog in the same context on the same day, the most ‘severe’ code assigned was retained for analysis.

#### 2.2.2 Post-adoption behaviour

On the point of adoption, adopters were asked to participate in a study evaluating the predictive accuracy of the shelter’s behavioural assessment and were given full details of the procedure (consent form provided in the Supplementary Materials). Consenting adopters each received two phone calls from a canine behaviourist at the shelter who surveyed them about their dog’s behaviour using a standardised set of questions (see Supplementary Materials). The majority of phone calls were made by one behaviourist although no records were kept on which behaviourist made the calls. The study plan was to record behaviour using telephone interviews at approximately 2-3 weeks and 5-6 weeks after adoption but the actual days after adoption were more variable. The average number of days after adoption for the first phone call was 19.8 (sd = 4.9; range = 7 to 51), and 32.7 days (sd = 11.1; range = 14 to 72) for the second call, with an average of 20.7 (sd = 5.6; range = 4 to 42) days between first and second interviews for those dogs with two completed surveys. Only 150 dogs (62%) had two fully complete surveys. One dog was returned after two post-adoption surveys and was subsequently adopted and received two surveys from their second owner; we randomly chose the first set of surveys on this dog for inclusion in the analysis. A different dog had been returned after the first survey and was subsequently adopted and received two surveys from their second owner; we chose the second owners’ surveys for this dog. Six additional dogs with only the first survey completed were subsequently returned to the shelter.

During the telephone survey, the first questions gathered information about an adopter’s dog ownership experience and the post-adoption environment. The remaining questions (Table 3) enquired about how the dog reacted in situations comparable to the eight behavioural observation contexts in the shelter assessment (Table 2). The questions were used as a guide during the post-adoption interviews, with the behaviourists providing more detailed descriptions as needed. The *In kennel* and *Out of kennel* contexts at the shelter were transformed to *In house* and *Out of house* (where house referred to the residence in which the dog lived with the adopter). The questions were open-ended, with the behaviourist encouraging adopters to describe the dog’s behaviour in each situation rather than label the behaviour. Subsequently, the behaviourist chose the behavioural code for each context that best described the behaviour. Adopters were allowed to respond ‘No opinion’, which was treated as a missing value. Phone calls were not recorded, precluding assessment of the reliability of behaviour coding during phone calls. Data were handled anonymously by the authors, who received only the dogs’ identification numbers to enable matching of the post-adoption reports with the shelter records.

### 2.3 Statistical analysis

#### 2.3.1 Theoretical approach

The green, amber and red codes from shelter and post-adoption time points were analysed using a joint or ‘shared parameter’ (e.g. Vonesh et al. 2006; Tseng et al. 2016) Bayesian hierarchical model, which accounted for two data complexities. First, we treated the missing data present in the shelter records (Table 2) and post-adoption surveys (Table 3) as non-ignorable (i.e. not random) because they likely depended on other variables. For example, the number of people unfamiliar to a dog decreases with time spent at the shelter or time after adoption, leading to potentially more missing records in the *Interactions with unfamiliar people* context. A dog’s overall behaviour could also impact missing values. For example, dogs with more green codes overall and who do not change their behaviour may not receive as many records due to a lack of priority relative to dogs who show more worrying or varying behaviour. The participation or attrition rate of adopters in the telephone surveys may also have depended on their dog’s behaviour post-adoption.

**Table 2:** *Observational contexts at the shelter with corresponding median and interquartile range (IQR; due to skewness) of the number of records per dog, and average percentage (standard deviation) of missing daily records per dog*.

| Context | Description | Median (IQR) | % Missing (SD) |
| --- | --- | --- | --- |
| Handling (HND) | Informal handling by people (e.g. stroking non-sensitive areas, touching or holding the collar, fitting a harness or lead). | 13 (14) | 40 (20) |
| In kennel (KNL) | Inside the kennel | 14 (14) | 37 (21) |
| Outside the kennel (OKNL) | Outside the kennel | 13 (14) | 42 (19) |
| Interactions with familiar people (FPL) | Interacts with familiar people (interacted with at least once before) outside the kennel who approach, make eye contact, speak to or attempt to make physical contact with the dog. | 13 (14) | 44 (18) |
| Interactions with unfamiliar people (UFPL) | Interacts with familiar people (never interacted with before) outside the kennel who approach, make eye contact, speak to or attempt to make physical contact with the dog. | 6 (5) | 75 (15) |
| Eating food (FOOD) | Eating food from a container, toy or while being hand-fed | 14 (13) | 39 (20) |
| Interactions with toys (TOYS) | Interacting with dog toys | 8 (13) | 58 (25) |
| Interactions with dogs (DOGS) | Meeting dogs outside the kennel during structured interactions and/or spontaneous meetings | 5 (4) | 76 (12) |

**Table 3:** *Post-adoption survey behaviour questions and response rates (number and percentage of owners responding to the question over two surveys)*.

| Context (abbreviation) | Survey question | *n* (%) with 2 surveys |
| --- | --- | --- |
| Handling (HND) | How has the dog behaved while being handled or restrained (informal, i.e. not in a veterinary context)? | 149 (61.8) |
| In house (HOUSE) | How has the dog behaved in the house? | 149 (61.8) |
| Outside of house (OTSD) | How has the dog behaved outside on walks? | 147 (61.0) |
| Interactions with familiar people (FPL) | How has the dog behaved when meeting familiar people? | 144 (59.8) |
| Interactions with unfamiliar people (UFPL) | How has the dog behaved when meeting unfamiliar people | 149 (61.8) |
| Eating food (FOOD) | How has the dog behaved with their food? | 150 (62.2) |
| Interactions with toys (TOYS) | How has the dog behaved with toys? | 149 (61.8) |
| Interactions with dogs (DOGS) | How has the dog behaved when meeting another dog for the first time? | 144 (59.8) |

A second complexity was implied by the high proportion (∼ 90%) of green codes, found in a previous analysis of data collected using the same shelter assessment (Goold and New-berry 2017a). Some proportion of the green codes may have been recorded due to other processes, such as staff members not observing the dog’s behaviour directly or forgetting how a dog behaved but inputting a green code anyway. Adopters may have also described their dog’s behaviour in terms consistent with green codes (i.e. not reporting any problems) during phone calls even if the dog’s behaviour might be more consistent with amber or red code behaviour. Thus, some green codes were potentially ‘inflated’, leading to a greater probability mass for green codes beyond that explained by the data generating processes of the behavioural scale alone. This ‘green code inflation’ assumption is similar to assumptions made when analysing count data with a high number of zero values (zero inflation; Lambert 1992) or ordinal data with a high probability mass for ‘I don’t know’ responses (Kelley and Anderson 2008) or presumed face-saving ‘Neither agree nor disagree’ responses (Bagozzi and Mukherjee 2012).

To account for the above complexities, we specified a custom mixture model for the probability of different codes (*c* = missing, green, amber, red = 0, 1, 2, 3) for case *i* (either the days after arriving at the shelter or the days after adoption), dog *j* and context *g*:

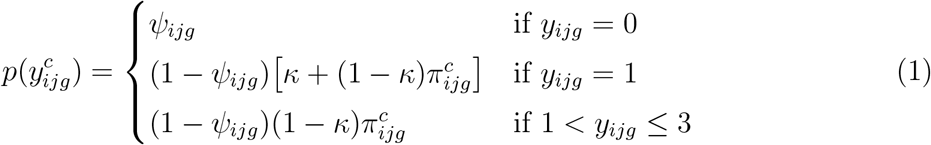

The parameter *ψ*_*ijg*_ is a ‘hurdle’ probability of missing data for each dog and context (see Kubinec 2019 for a similar approach to handling missing data), which is modelled as a Bernoulli trial from the data:

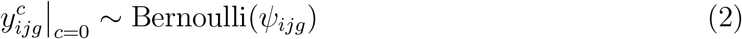

The complement, 1 − *ψ*_*ijg*_, is the probability of either a staff member choosing to input an observation in the shelter or an adopter participating in the telephone survey, respectively. The *κ* parameter is a mixing term that describes the probability of a green code being drawn from the inflation component. Consequently, 1 − *κ* is the probability of a code being non-inflated. A non-missing and non-inflated code of category *c* (for a specific case, dog and context) occurs with probability 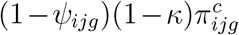, where 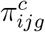 is defined using a cumulative ordinal probit model:

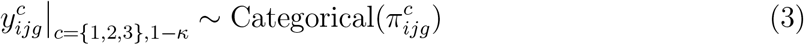

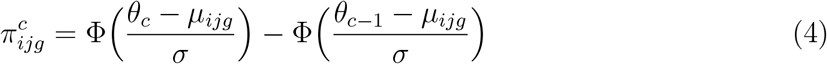

Equation 3 describes the non-missing and non-inflated code as categorically distributed with probability 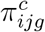, which is defined in equation 4 as the cumulative area under a latent standard normal distribution (where Φ is the standard normal cumulative distribution function) between threshold parameters *θ*_*c*_ and *θ*_*c*−1_ (***θ*** = *θ*_0_, *θ*_1_, *θ*_2_, *θ*_3_). The probability of the first and last threshold parameters (i.e. Φ(*θ*_0_), Φ(*θ*_3_)) were set to 0 and 1, respectively. To estimate the mean (*μ*_*ijk*_) and standard deviation (*σ*) on the scale of the ordinal data, we fixed *θ*_1_ = 1.5 and *θ*_2_ = 2.5 (see Kruschke 2014).

#### 2.3.2 Joint modelling

The data from the shelter and post-adoption time periods were each analysed by the model described above, which we describe as two distinct ‘sub-models’, both predicting the probability of missing data (*ψ*_*ijg*_ in equation 2 using a logit link function) and the predicted mean of the latent behavioural scale scores (*μ_ijg_* in equation 4 using the identity link function). Each sub-model included the dog-, context-, and dog × context-varying intercept and slope parameters (for the number of days after arrival at the shelter or days after adoption, respectively), which represent personality (intercept parameters) and plasticity (slope parameters), respectively. The joint structure of our model accounts for correlations between these parameters across the shelter and post-adoption sub-models. For the ordinal probit regressions (equation 4), we included random intercepts and slope terms for dogs, contexts and the interaction between dogs and contexts (1, 928 unique dog × contexts combinations). Dog- and context-varying effects capture additive variation due to dogs and contexts, respectively, while the dog × context interaction describes the non-additive, unique variation attributed to specific dogs in specific contexts. For the missing data regressions, we included only random intercepts. For each random effect term (dogs, contexts, and dogs × contexts), this led to a 6 x 6 covariance matrix capturing the relationship between random intercepts and slopes for the shelter and post-adoption behaviours. We summarised the variation across dogs, contexts and dogs within contexts by calculating behavioural repeatability, the proportion of model variance explained by the variance of the random intercept parameters (see the Supplementary Materials).

#### 2.3.3 Predictor variables

For the shelter sub-model, we included days after arrival at the shelter as an observation-level predictor and the variables from Table 1 as dog-level predictors, excluding the total number of observations (which was nearly perfectly collinear with length of stay) and rehoming centre (which was not of interest *a priori*). The post-adoption sub-model included days after adoption as an observation-level predictor and the same shelter demographic variables as in the shelter sub-model as retrospective dog-level predictors, as well as the number of post-adoption surveys about each dog. Sum-to-zero coding was used for categorical predictors. Dog-level metric predictors were mean-centred and standardised by 1 standard deviation. The number of days after arrival at the shelter was centered around the average length of stay at the shelter (32.1 days; Table 1) and scaled by 1 standard deviation (73 days). The number of days after adoption was centered around the average of each dog’s mean number of days after adoption (30.1 days) and scaled by its standard deviation (∼ 11.6 days). One dog had missing neuter status data, two dogs were missing weight data and two dogs were missing age at departure data. Due to the small amount of missing data, we imputed the missing neuter status data point with the most frequent category (neutered on site; Table 1), and mean imputed the missing weight and age data points.

#### 2.3.4 Estimation & inference

All data cleaning and post-processing was conducted in R version 3.6.1 (R Core Team 2019). We fit our model using Bayesian estimation in the Stan programming language (Stan Development Team 2018) using the terminal interface CmdStan version 2.23.0 (Stan Development Team et al. 2018). Stan employs Hamiltonian Monte Carlo, a type of Markov chain Monte Carlo (MCMC) sampling algorithm, to sample from the posterior distribution. After ensuring adequate mixing of multiple chains from the model ran on smaller sub-sets of the data, we took 10,000 iterations from the posterior distribution using 2 MCMC chains (1000 warmup and 5000 sampling). We summarise parameters by their means and 95% highest density intervals (HDIs). For the ordinal data, we make predictions using both the latent metric scale (parameters denoted by *β*) and the corresponding posterior probabilities of each ordinal category (probabilities denoted by *π*). When evaluating population-level parameters, we highlight discrepancies, where appropriate, between the estimates for average dogs and contexts, and estimates that incorporate uncertainty due to variation across dogs and contexts. The former represent estimates made at the mean of the random effects, while the latter are made by marginalising over the random-effect distributions and are particularly important for making accurate future predictions in hierarchical models (e.g. IntHout et al. 2016; Wang and Lee 2019).

### 2.4 Ethical statement

Approval for the processing of personal data was obtained from the Norwegian Social Science Data Services (approval number 47080). Names and addresses of the participating new owners and their dogs were anonymised before data were passed to the authors.

### 2.5 Data & code accessibility

We provide complete mathematical details of the above model, supplementary files, the data, the Stan code, and the R code to reproduce the results reported here at https://github.com/cmgoold/goold-newberry-lba. Due to the non-standard statistical model, we also provide R and Stan code to simulate data and fit the model to recover parameter values for model validation.

## 3 Results

### 3.1 Average behaviour

Green codes accounted for more than 95% of the observations in the raw shelter and post-adoption datasets. For an average dog in an average context (i.e. at the mean of the random effects, where all dog-level predictors equalled 0), there were no credible differences between the probabilities of green, amber and red codes at the shelter (at approximately 32.1 days after arrival) and post-adoption (at 30.1 days post-adoption; probability differences: *π*_green_ = 0.00, 95% HDI = [-0.05, 0.06]; *π*_amber_ = 0.00, 95% HDI: [-0.06, 0.05]; *π*_red_ = 0.00, 95% HDI: [0.00, 0.00]). For an average dog in an average context, behaviour tended to improve with every additional week at the shelter (*β* = −0.08; 95% HDI: [-0.14, -0.02]), although due to ceiling effects the practical increase in green codes was minimal (from around 98% on arrival day to 99% on day 30). There was no clear relationship between days after adoption and post-adoption behaviour (*β* = 0.01; 95% HDI: [-0.21, 0.24]). A dog’s average probability of missing data at the shelter (*π*_missing_) was 0.54 (95% HDI: [0.37, 0.73]), and the odds of missing codes increased with every additional week after arrival (odds: 1.04; 95% HDI: [1.03, 1.05]). The odds of missing data post-adoption increased sharply with every additional week after adoption (odds: 11.62; 95% HDI: [8.90, 14.00]), although this was mainly a between-survey phenomenon driven by dogs with only 1 completed survey, whereas within-survey missing data for dogs with two completed surveys were highly infrequent (approximately zero).

The probability of green-code inflation was *κ* = 0.04 (95% HDI: [0.00, 0.09]) at the shelter and *κ* = 0.20 (95% HDI: [0.03, 0.36]) post-adoption.

### 3.2 Variation across dogs and contexts

When accounting for variation across dogs and contexts, there was substantially more uncertainty in the model estimates both at the shelter (Fig. 1) and post-adoption (Fig. 2). At the shelter, the *Interactions with dogs* context had the highest probability of red codes (*π*_red_ = 0.04; 95% HDI: [0.02, 0.06]; Supplementary Fig. S1a), and dogs changed their behaviour the most in the *Handling* context (*β* = −0.20 95% HDI: [-0.28, -0.14]; Supplementary Fig. S1b). Post-adoption, the *Eating food* context had the highest probability of red codes (*π_red_* = 0.01; 95% HDI: [0.00, 0.05]; Supplementary Fig. S2a), although there were no strong differences between contexts in the average amount of behavioural change across days after adoption (Supplementary Fig. S2b). Missing codes were particularly prominent at the shelter for the *Interactions with unfamiliar people* context (*π_missing_* = 0.81; 95% HDI: [0.74, 0.89]).

**Figure 1:**
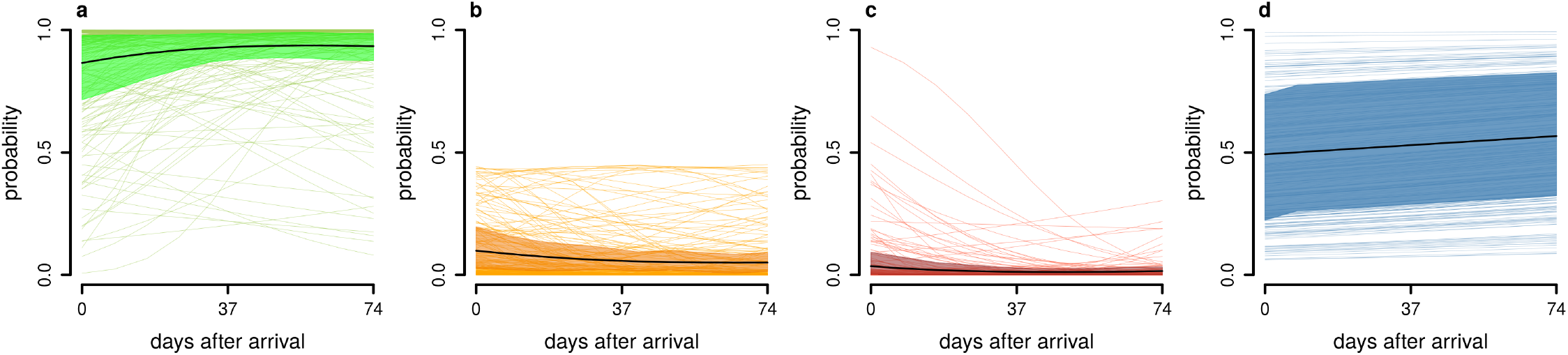
*Probability of different behaviour codes across days after arrival at the shelter (average length of stay + 1 standard deviation) accounting for variation in behaviour across dogs, contexts and dog × context; (a) green, (b) amber, (c) red, (d) missing. Thick black lines show posterior mean estimates, coloured bands show the 95% highest density interval of the mean, and thin coloured lines (n = 241) show one random sample from one randomly-chosen context from each dog’s posterior distribution*.

**Figure 2:**
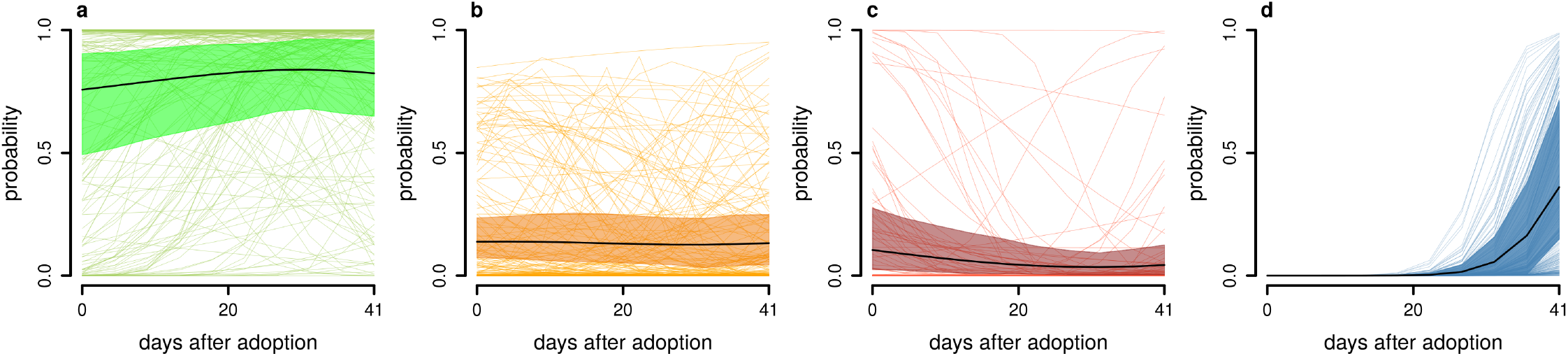
*Probability of different behaviour codes across days after adoption (average days after adoption interview + 1 standard deviation) accounting for variation across dogs, contexts and dog × context behaviour; (a) green, (b) amber, (c) red, (d) missing. Thick black lines show posterior mean estimates, coloured bands show the 95% highest density interval of the mean, and thin coloured lines (n = 241) show one random sample from one randomly-chosen context from each dog’s posterior distribution*.

Behavioural repeatability was highest across contexts at the shelter and for dogs within contexts in the post-adoption surveys (Fig. 3). Repeatability for dogs within contexts was approximately 20% higher post-adoption than at the shelter (*R_diff_* = −0.23; 95% HDI: [-0.41, -0.05]) but was more uncertain, with the standard deviation of the dog × context-varying intercepts being 1.40 (95% HDI: [1.10, 1.72]) times higher post-adoption than at the shelter. The standard deviation of the dog × context slope parameters was considerably larger at the shelter (describing behaviour over days after arrival) than post-adoption (describing behaviour over days after adoption), but had substantial uncertainty (ratio of standard deviations: 45.54; 95% HDI: [1.61, 96.43]).

**Figure 3:**
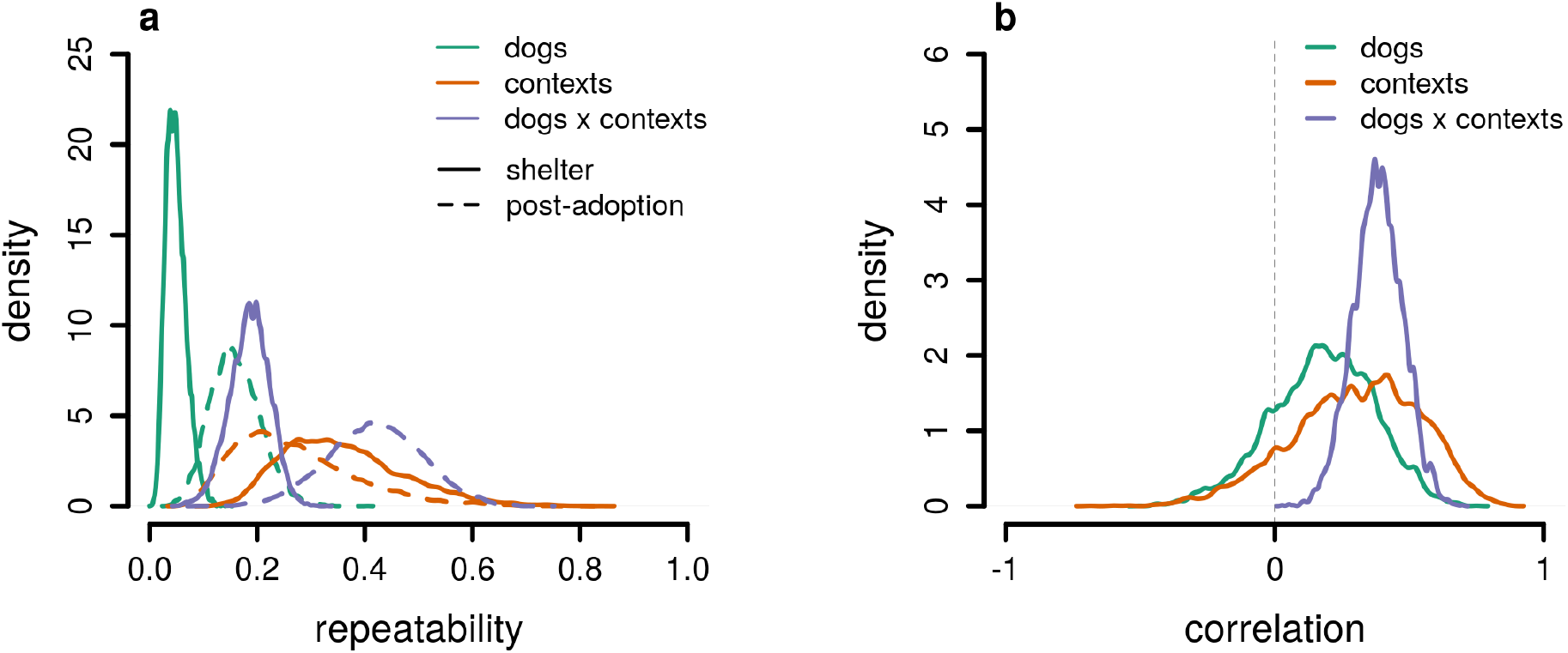
*Behavioural differences between dogs, contexts and dogs within contexts at the shelter and post-adoption; (a) proportions of total behavioural variance (repeatability) explained by dog-, context- and dog × context-varying intercepts for shelter and post-adoption time periods, and (b) correlations between shelter and post-adoption time periods for dog-, context- and dog × context-varying intercepts (Supplementary Table S1)*.

**Figure 4:**
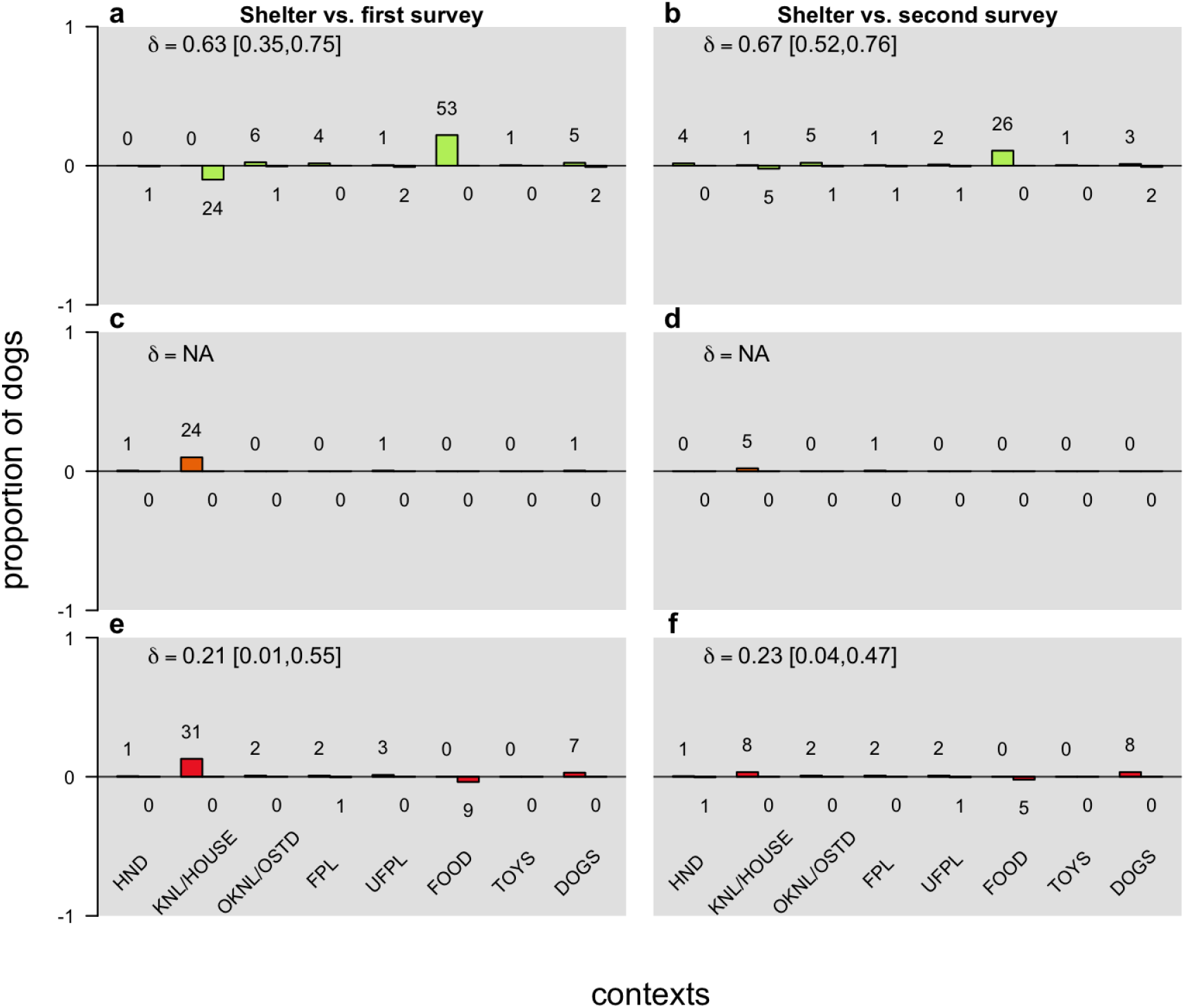
*The proportion (bars) and number of dogs (text above each bar) who had significantly different probabilities of green (a-b), amber (c-d), and red (e-f) codes within contexts at the shelter compared to behaviour reported at the first (a, c, e) and second (b, d, f) post-adoption phone calls. Positive bars illustrate the proportion of dogs with higher code probabilities at the shelter compared to post-adoption; negative bars illustrate the proportion of dogs with lower probabilities at the shelter compared to post-adoption. A dog was included in the count if the 95% highest density interval of the differences in code probabilities for a dog did not include zero. The mean (δ = [0, 1]) and 95% highest density interval absolute probability difference is shown above each plot, except for panels c and d where there were too few dogs for its calculation*.

**Figure 5:**
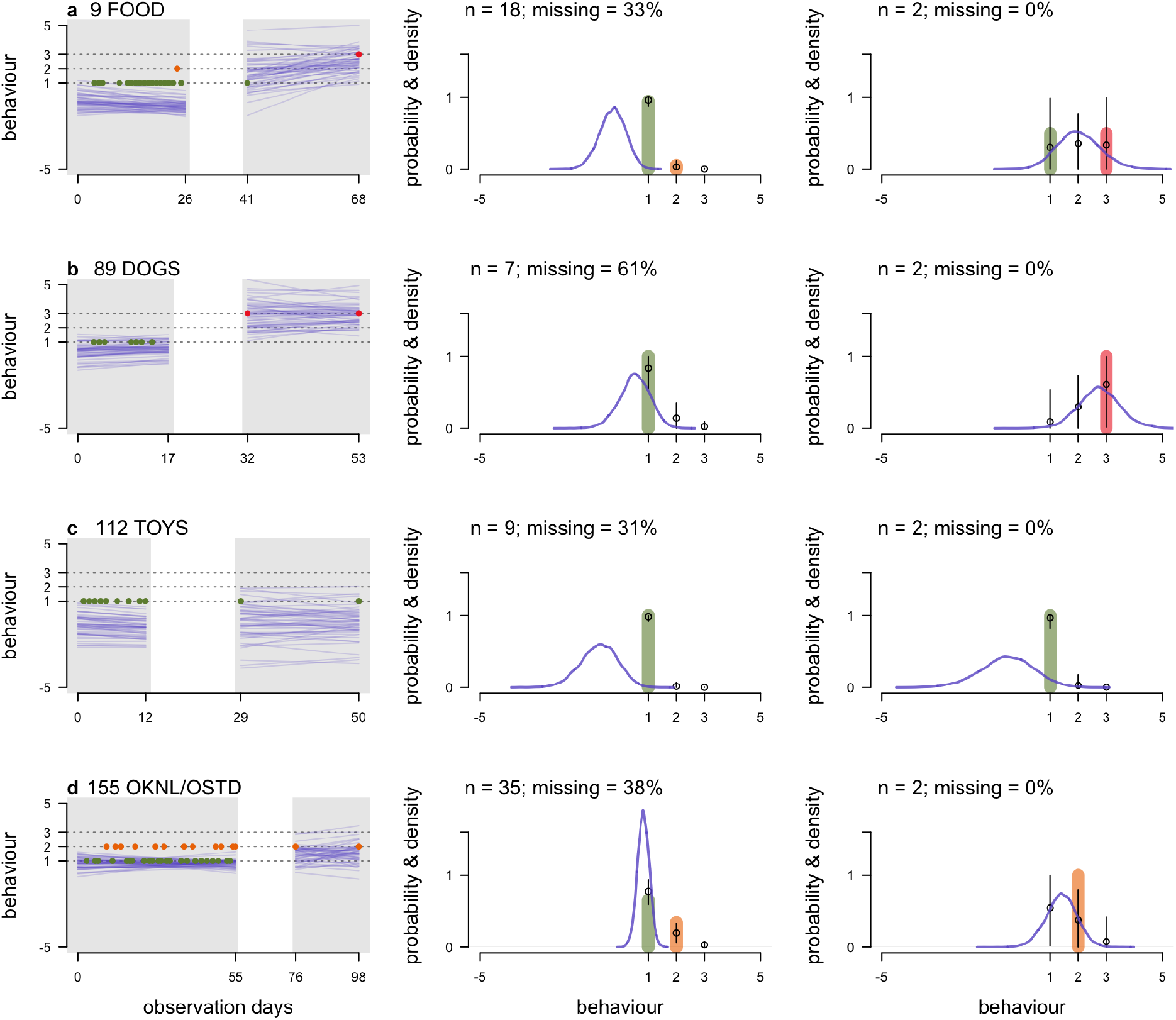
*Posterior predictions for 4 specific dogs and contexts (a-d; anonymous dog ID number and context abbreviation shown above first panel of each plot; OKNL/OSTD = outside of kennel/house, FOOD = eating food, TOYS/DOGS = interactions with toys/dogs; see Supplementary Materials for more examples). Left panels show behaviour over days after arrival at the shelter (first gray area) and behaviour over days after adoption (second gray area), where the coloured dots denote the raw ordinal data (green=1, amber=2, red=3, where red denotes the most undesirable behaviour) and blue lines 50 samples from the posterior distribution on the latent behavioural scale. Middle and right panels show the raw proportions of green, amber and red codes (coloured bars) at the shelter (middle) and post-adoption (right), with the number of observations and percent missing data displayed above; black dots and vertical line segments denote the mean and 95% highest density interval of the predicted probabilities of each code. Blue density curves show the estimated latent scale generating the ordinal codes for each dog*.

### 3.3 Random effect correlations

The dog × context-varying intercept parameters at the shelter and post-adoption had a moderate positive correlation (*ρ* = 0.38; 95% HDI: [0.20, 0.55]; Fig. 3b), whereas the correlations between dog-varying and context-varying intercepts were largely positive but their 95% HDIs spanned zero (Supplementary Table S1). Correlations between the random-slope parameters at the shelter and post-adoption included zero. At the shelter, the dog × context intercept and slope parameters were positively correlated (*ρ* = 0.26; 95% HDI: [0.02, 0.47]), meaning that dogs with higher values on the latent behavioural scale (higher changes of red codes) changed their behaviour less over time or, in some rare cases, had positive slope parameters meaning they had increased chances of exhibiting amber and red codes over time. Dogs with more positive slope parameters across contexts at the shelter had slightly higher chances of missing data at the shelter (*ρ* = 0.28; 95% HDI: [0.01, 0.54]). Dogs with fewer missing data on average post-adoption also had larger intercept parameters (*ρ* = −0.38; 95% HDI: [-0.71, -0.04]) and more positive slopes (*ρ* = −0.36; 95% HDI: [-0.69,-0.02]) at the shelter, suggesting that adopters of dogs with overall higher chances of amber and red codes at the shelter had better response rates in the post-adoption phone calls.

We note that average post-adoption behaviour had a mostly positive correlation with the amount of missing data post-adoption, although the 95% HDI was wide and included zero (*ρ* = 0.25; 95% HDI: [-0.07, 0.53]).

### 3.4 Individual-level prediction of post-adoption behaviour

Behaviour was not static across shelter and post-adoption time points. A median of 48.50 (inter-quartile range: 32.75) and 39 (inter-quartile range: 66.75) dogs who received at least one amber and red code, respectively, within contexts at the shelter did not show amber or red codes post-adoption for the same contexts. Similarly, a median of 29.50 (inter-quartile range: 56.75) and 13 (inter-quartile range: 65.25) dogs who only showed green or amber codes at the shelter within contexts, received red codes in the post-adoption reports. As mentioned earlier, these diagnostic values only provide a crude estimate of behavioural differences between the shelter and post-adoption time periods, ignoring unbalanced repeated measures, measurement error and missing data. They further ignore each dog’s latent probabilities of showing green, amber and red behaviours.

Therefore, we compared the posterior probabilities of green, amber and red codes on each dog’s mean day after arrival at the shelter with the equivalent codes’ predicted probabilities on the day of the first and second telephone survey (where dogs with only the first survey had their second survey predictions estimated by the model). We summarised the resulting differences (*δ*) in code probabilities by counting the number and proportion of dogs whose 95% HDIs of the differences did not contain zero (Fig. 4). In general, the number of dogs who had significantly different probabilities of green, amber or red codes post-adoption than at the shelter was low (mean of estimates: 2.83; median: 0; sd: 7.51; range: 0 to 53). There were fewer overall differences between behaviour reported at the second surveys versus at the shelter than between the first survey versus at the shelter, likely due to fewer second surveys being available resulting in increased uncertainty in the predictions. Notably, approximately 10% of dogs (n = 24) had lower probabilities of green codes, and 10% (n = 24) and 13% (n = 31) of dogs had higher chances of amber and red codes, respectively, in the *In kennel* context at the shelter than the corresponding *In house* context at the first survey, although this discrepancy had largely disappeared by the second survey phone call. Around 20% of dogs (n = 53) had higher probabilities of green codes in the *Eating food* context at the shelter compared to the first survey, dropping to approximately 11% (n = 26) at the second survey. A smaller number of dogs (9 and 5 at first and second surveys, respectively) had increased chances of red codes post-adoption in the *Eating food* context.

### 3.5 Dog-level predictors

Length of stay at the shelter had a positive (higher latent scale scores) but weak relationship with dogs’ average behaviour at the shelter (*β* = 0.02; 95% HDI: [0.01, 0.03]) and post-adoption (*β* = 0.02; 95% HDI: [0.00, 0.03]), meaning that dogs that remained longer at the shelter had more amber and red codes. Dogs neutered before arrival at the shelter had higher latent scale scores on average than intact dogs (*β*_[no−yes]_ = −0.38; 95% HDI: [-0.73, -0.07]), with dogs neutered at the shelter and dogs of unknown neuter status having intermediate scores. Weight had a negative relationship (green codes increased with weight) with average behaviour post-adoption (*β* = −0.15, 95% HDI: [-0.26, -0.05]). Approximate age when leaving the shelter, source (gift/stray/return) and sex of dog were not significantly predictive of personality or plasticity at the shelter or post-adoption. None of the predictors significantly predicted the probability of missing data.

## 4 Discussion

This study provides the first comprehensive analysis for predicting shelter dog behaviour post-adoption from a longitudinal behavioural assessment. We applied a novel Bayesian hierarchical model to account for both shelter and post-adoption behaviour, missing data, and measurement error within a single statistical framework, which allowed individual-level inferences regarding how dogs would behave in specific contexts at the shelter and in their new home. We found few differences between how likely dogs were to show green, amber and red-coded behaviour on average at the shelter, and their corresponding chances of showing the same behavioural codes post-adoption. Moreover, in contrast to some previous studies reporting high numbers of behavioural problems in shelter dogs (e.g. Gates et al. 2018), the behavioural reports in this study were largely dominated by green codes (i.e. the most favourable behaviour). This does not imply that dogs behaved exactly the same post-adoption as they did in the shelter, however. Due to the varying amounts of data available per dog per context, there was considerable uncertainty in individual-level, latent probabilities of green, amber and red codes, particularly in the post-adoption reports where dogs had a maximum of two records per context. This makes it plausible, for example, that a dog who only displayed green or amber codes at the shelter but showed red code behaviour post-adoption could have statistically similar latent probabilities of different behaviours at the shelter and post-adoption. Far from simply being categorised as a ‘false negative’, longitudinal assessment and our inferential framework provides a richer, probabilistic assessment of individual dog behaviour on which to make more holistic assessments.

The hierarchical structure of our data allowed comparing variation in the behavioural records across dogs, across contexts, and across specific dogs × contexts combinations. At the shelter, the largest portion of behavioural variance was explained by variation across contexts, while variation was greatest between dogs within contexts in the post-adoption surveys (Fig. 3a). Contexts at the shelter were therefore more distinct from each other than post-adoption, which is evident in Supplementary Figures S1a and S2a where the estimated intercepts for the shelter contexts are more varied than for the post-adoption contexts. For instance, the *Interactions with dogs* context had much higher probabilities of red codes at the shelter than any other context. The greater uncertainty of the post-adoption context estimates also reduced their variation. The correlations between shelter and post-adoption behaviour were only credibly larger than zero for the dog × context-varying intercept parameters (Fig. 3b). Neither how individual dogs behaved across contexts on average, nor average dog behaviour within contexts, were particularly illuminating for predicting behaviour post-adoption. Repeatability was also around 20% higher post-adoption than in the shelter for the dog × context behaviour, driven by greater heterogeneity in the post-adoption behaviour estimates than the shelter reports.

The positive correlation between dog behaviour within contexts at the shelter and post-adoption informs us about their linear association, but the practical implications of this positive correlation are not easy to discern. One option is to compute rates of false positives and negatives (e.g. Marder et al. 2013) or positive and negative predictive values (e.g. Pa-tronek and Bradley 2016) to understand an assesment’s predictive ‘ability’ (Patronek et al. 2019). This medical diagnostic approach lends itself easily to scenarios whereby behavioural testing is conducted once at the shelter and once post-adoption, and for test protocols where the outcome measure is a binary pass or fail result. However, moving away from battery-style behavioural testing at shelters necessitates moving away from labelling dogs as either problematic or not problematic, or aggressive or non-aggressive, from limited behavioural information. Such sharp behavioural categorisations are at odds with the rationale for longitudinal assessments (ASPCA 2018; Rayment et al. 2015) that encourage forming a holistic profile of a dog’s behaviour over different contexts, rather than focusing on any one single behaviour. Instead, we leveraged the posterior predictions of our model to determine how similar dog behaviour was across the post-adoption behaviour reports compared to each dog’s behaviour on average at the shelter. Hierarchical models are excellently suited to this task because they balance the information from each dog with what is typical across the whole sample. This ‘partial pooling’ of information across dogs and contexts makes better predictions than non-hierarchical models or raw data because it balances over- or under-fitting to the data (e.g. Gelman and Hill 2007; McElreath and Koster 2014).

To pick a few representative examples (see Supplementary Materials for more examples), Fig. 5 illustrates the posterior predictions for four dogs, each in a different context, with the raw data for comparison. On the estimated latent scale generating the ordinal data, predictions for dogs’ post-adoption behaviour (indicated by the sample of regression lines in the left-hand plots) tended to be more uncertain than their in-shelter predictions, where more data was available overall (e.g. compare shelter and post-adoption results in Fig. 5a). The impact of the amount of data available can also be seen by inspecting the overall probabilities of different code colours, shown in the middle and right panels of each plot, where behaviour of dogs with fewer data points was more uncertain (e.g. the 95% HDIs in the middle panel of Fig. 5b). What occurs on average in certain contexts also affects the individual-level predictions. For instance, the dog in Fig. 5b received no red codes at the shelter in the *Interactions with dogs* context, but received red codes post-adoption. The overall increase in this dog’s probability of red codes in the latter context compared to the shelter was 56%, but with a 95% highest density interval that spans from 100% to-1%, meaning there was a small chance that red codes were just as, or more, likely at the shelter than post-adoption. While this might seem surprising, it occurs because red codes were most probable at the shelter in the *Interactions with dogs* context overall, so even dogs with no red codes in this context had an increased chance of red codes compared to other contexts. The *Interactions with dogs* context also had one of the highest amounts of missing data (e.g. dog 89 in Fig. 5b has only 7 observations, with 61% missing data), and thus the predictions are more uncertain overall. By contrast, green codes were particularly dominant in the *Interactions with toys* context, and the amount of missing data was lower than average (Supplementary Fig. S1b), so dogs showing green codes in this context at the shelter and post-adoption had much tighter posterior estimates (e.g. see the predicted probabilities for the dog in Fig. 5c).

For most contexts, the number of dogs likely to show differences in probabilities of green, amber and red codes at the shelter compared to post-adoption was low (Fig. 4). This supports the ability of the longitudinal assessment to predict post-adoption behaviour, although we emphasise that this good predictive ability is in part due to the appropriate representation of uncertainty in the estimates. The greatest discrepancies between shelter and post-adoption behaviour appeared in the *In kennel/In house* and *Eating food* contexts. Between 10 and 15% of dogs in the former context were predicted to show more green codes, and fewer amber and red codes, at the first post-adoption phone call than at the shelter (Fig. 4). This is perhaps unsurprising as the *In kennel* and *In house* contexts were the least comparable contexts across shelter and post-adoption periods, and a kennel would be expected to be a less stressful environment than the adoptive home. In contrast, around 20% of dogs at the first post-adoption survey, and around 11% at the second survey, showed lower chances of green codes than at the shelter in how they behaved around their food. The latter result is partly consistent with previous research questioning whether reasonable predictions about how dogs behave around food in the adopted home can be made from their behaviour at the shelter (e.g. Marder et al. 2013; Mohan-Gibbons et al. 2012; Clay et al. 2020a). We also found that, post-adoption, the probability of red codes was highest in the *Eating food* context. However, the latter probability was still very low (∼ 1%), and apart from nine dogs at the first survey and five dogs at the second, we found no evidence for increased probabilities of red codes post-adoption for behaviour around the food bowl. Thus, repeated observational data collected on how dogs behave around their food bowl at the shelter may provide a more valid (in the predictive ability sense) indicator of post-adoption behaviour, even in this historically difficult-to-infer context.

Longitudinal behavioural assessments and adopter-reported behaviour are at risk of both missing data and measurement error, which are unlikely to be avoided entirely. Instead, understanding when and how they emerge is key to accounting for any deleterious effects they have on behavioural predictions. We found that dogs’ odds of missing data at the shelter were positively correlated with the behavioural plasticity across days after arrival. Given that the average coefficient describing behavioural change at the shelter was negative (indicating an improvement in behaviour through time), and that most dogs had largely negative individual slope estimates (i.e. deterioration in behaviour was rare), the latter correlation means that dogs who had shallower slopes (i.e. less plastic) were more likely to have missing data on average at the shelter. Importantly, there was no evidence to suggest that dogs with less favourable behaviour at the shelter on average were more likely to have missing data (Supplementary Table S1). We further found that owner response rates in the post-adoption surveys correlated more clearly with dog behaviour reported in the shelter than behaviour reported post-adoption: dogs with higher latent scale averages at the shelter, as well as dogs who changed less over time at the shelter, had lower chances of missing data post-adoption. We refrain from hypothesising-after-the-results-are-known (i.e. HARKing; Kerr 1998) on the causal mechanisms driving these correlations, but do emphasise the importance of understanding how response patterns (from shelter staff and new owners) are impacted by dog behaviour for the efficacy of longitudinal behaviour assessments. We also avoid speculating about reasons for the associations found between the dogs’ behaviour and length of stay at the shelter, body weight (which is related to breed type) and neuter status, but note that these variables explained some of the variance, supporting their inclusion in models of shelter dog behaviour. Age, sourcing from different shelter facilities and sex of dog did not contribute significantly to behaviour when assessed on the shelter’s ‘desirability for adoption’ scale, although these factors could contribute to more specific behavioural differences and to interactions that were not evaluated.

The post-adoption behaviour reports had larger measurement error estimates than the shelter reports. Like estimates of zero-inflation in count data (Lambert 1992), we estimated ‘green code inflation’, that is, the proportion of green codes reported due to processes different from the process generating the behavioural data. In reality, the inflation component is a place-holder for more complex mechanisms, which could include shelter staff not observing a dog’s behaviour but inputting a green code anyway, or adopters reporting behaviours consistent with green codes, even when the real behaviour was less favourable. Green codes had approximately a 20% chance of being drawn from the inflation component of the model post-adoption, although there was large uncertainty (95% HDI: [3, 36]). By comparison, green code inflation was estimated at only 4% at the shelter. While we cannot disentangle the adopters’ own behavioural reports from how the shelter behaviourists interpreted the behavioural descriptions from the adopters, this implies that systematic error in the adopter-reported behavioural reports is important to quantify further. Unfortunately, no previous studies have assessed patterns of measurement error in shelter or post-adoption behaviour.

The modelling approach in this study is provided completely open source, and we encourage future studies predicting shelter dog behaviour to consider principled statistical frameworks such as hierarchical Bayesian methods. For example, if a larger sample of data was available, this study could have trained the model on a sub-sample of the data and made out-of-sample predictions about post-adoption for the remaining data. Moreover, due to the cumulative nature of Bayesian inference (McElreath 2020), the modelling approach here could be applied in real-time at shelters to provide predictions for shelter dog behaviour both while at the shelter and post-adoption. The relative probabilities of dogs showing different types of behaviours could be integrated with qualitative information on each individual dog (e.g. pre-surrending surveys; ASPCA 2018) and expert knowledge to form an informed adoption decision and find a suitable new home.

## 5 Conclusion

Predicting the future behaviour of shelter dogs is a difficult task. Longitudinal assessments have been proposed as an alternative to battery-style, standardised testing, but research is still scarce on their implementation and efficacy. We have provided the first comprehensive analysis of a longitudinal assessment to predict the behaviour of shelter dogs post-adoption using a novel application of Bayesian hierarchical modelling. While dog behaviour was not static, few dogs showed marked changes in their latent behavioural profiles at the shelter and post-adoption, supporting the use of longitudinal assessments and our inferential framework to inform adoption decisions alongside additional information and expert judgement. We have, further, demonstrated links between dog behaviour, missing data and measurement error, which should be accounted for in future analyses of longitudinal assessment data.

## References

ASPCA (2018). Position statement on shelter dog behavior assessments. https://www.aspca.org/about-us/aspca-policy-and-position-statements/position-statement-shelter-dog-behavior-assessments. Online; accessed 13-May-2020.

Bagozzi, B. E. and B. Mukherjee (2012). “A mixture model for middle category inflation in ordered survey responses”. In: Political Analysis 20.3, pp. 369–386.

Bollen, K. S. and J. Horowitz (2008). “Behavioral evaluation and demographic information in the assessment of aggressiveness in shelter dogs”. In: Applied Animal Behaviour Science 112.1–2, pp. 120–135.

Borsboom, D., G. J. Mellenbergh, and J. Van Heerden (2004). “The concept of validity.” In: Psychological Review 111.4, p. 1061.

Borsboom, D. et al. (2009). “The end of construct validity.” In: The Concept of Validity: Revisions, New Directions and Applications, Oct, 2008. IAP Information Age Publishing.

Christensen, E. et al. (2007). “Aggressive behavior in adopted dogs that passed a tempera-ment test”. In: Applied Animal Behaviour Science 106.1–3, pp. 85–95.

Clay, L. et al. (2020a). “Do behaviour assessments in a shelter predict the behaviour of dogs post-adoption?” In: Animals 10.7, p. 1225.

Clay, L. et al. (2020b). “In Defense of Canine Behavioral Assessments in Shelters: outlining Their Positive Applications”. In: Journal of Veterinary Behavior.

Cleasby, I. R., S. Nakagawa, and H. Schielzeth (2015). “Quantifying the predictability of behaviour: statistical approaches for the study of between-individual variation in the within-individual variance”. In: Methods in Ecology and Evolution 6.1, pp. 27–37.

Dingemanse, N. J. et al. (2010). “Behavioural reaction norms: animal personality meets individual plasticity”. In: Trends in Ecology & Evolution 25.2, pp. 81–89.

Gates, M. C. et al. (2018). “Post-adoption problem behaviours in adolescent and adult dogs rehomed through a New Zealand animal shelter”. In: Animals 8.6, p. 93.

Gelman, A. and J. Hill (2007). Data Analysis using Regression and Multilevel/Hierarchical Models. Vol. 1. Cambridge University Press New York, NY, USA.

Goold, C. and R. C. Newberry (2017a). “Modelling personality, plasticity and predictability in shelter dogs”. In: Royal Society Open Science 4.9, p. 170618.

Goold, C. M. and R. C. Newberry (2017b). “Aggressiveness as a latent personality trait of domestic dogs: testing measurement invariance and local independence”. In: PLoS One.

Hofstadter, A. (1951). “Explanation and necessity”. In: Philosophy and Phenomenological Research 11.3, pp. 339–347.

IntHout, J. et al. (2016). “Plea for routinely presenting prediction intervals in meta-analysis”. In: BMJ Open 6.7, e010247.

Kelley, M. E. and S. J. Anderson (2008). “Zero inflation in ordinal data: incorporating susceptibility to response through the use of a mixture model”. In: Statistics in Medicine 27.18, pp. 3674–3688.

Kerr, N. L. (1998). “HARKing: Hypothesizing after the results are known”. In: Personality and Social Psychology Review 2.3, pp. 196–217.

Kruschke, J. (2014). Doing Bayesian Data Analysis: A tutorial with R, JAGS, and Stan. Academic Press.

Kubinec, R. (2019). “Generalized Ideal Point Models for Time-Varying and Missing-Data Inference”. In: OSF Preprint.

Lambert, D. (1992). “Zero-inflated Poisson regression, with an application to defects in manufacturing”. In: Technometrics 34.1, pp. 1–14.

Marder, A. R. et al. (2013). “Food-related aggression in shelter dogs: a comparison of behavior identified by a behavior evaluation in the shelter and owner reports after adoption”. In: Applied Animal Behaviour Science 148.1–2, pp. 150–156.

Marston, L. C. and P. C. Bennett (2003). “Reforging the bond—towards successful canine adoption”. In: Applied Animal Behaviour Science 83.3, pp. 227–245.

Maul, A., D. T. Irribarra, and M. Wilson (2016). “On the philosophical foundations of psychological measurement”. In: Measurement 79, pp. 311–320.

McElreath, R. (2020). Statistical Rethinking: A Bayesian Course with Examples in R and Stan. CRC press.

McElreath, R. and J. Koster (2014). “Using multilevel models to estimate variation in foraging returns”. In: Human Nature 25.1, pp. 100–120.

Mohan-Gibbons, H., E. Weiss, and M. Slater (2012). “Preliminary investigation of food guarding behavior in shelter dogs in the United States”. In: Animals 2.3, pp. 331–346.

Mohan-Gibbons, H. et al. (2018). “The impact of excluding food guarding from a standardized behavioral canine assessment in animal shelters”. In: Animals 8.2, p. 27.

Mornement, K. et al. (2009). “Reliability, validity and feasibility of existing tests of canine behaviour”. In: Proceedings of AIAM Annual Conference on Urban Animal Management, pp. 11–18.

Mornement, K. M. et al. (2010). “A review of behavioral assessment protocols used by Australian animal shelters to determine the adoption suitability of dogs”. In: Journal of Applied Animal Welfare Science 13.4, pp. 314–329.

Mornement, K. M. et al. (2015). “Evaluation of the predictive validity of the Behavioural Assessment for Re-homing K9’s (BARK) protocol and owner satisfaction with adopted dogs”. In: Applied Animal Behaviour Science 167, pp. 35–42.

Patronek, G. J. and J. Bradley (2016). “No better than flipping a coin: reconsidering canine behavior evaluations in animal shelters”. In: Journal of Veterinary Behavior 15, pp. 66–77.

Patronek, G. J., J. Bradley, and E. Arps (2019). “What is the evidence for reliability and validity of behavior evaluations for shelter dogs? A prequel to “No better than flipping a coin””. In: Journal of Veterinary Behavior.

Poulsen, A., A. Lisle, and C. Phillips (2010). “An evaluation of a behaviour assessment to determine the suitability of shelter dogs for rehoming”. In: Veterinary Medicine International 2010.

R Core Team (2019). R: A Language and Environment for Statistical Computing. R Foundation for Statistical Computing. Vienna, Austria. url: https://www.R-project.org/.

Rayment, D. J. et al. (2015). “Applied personality assessment in domestic dogs: limitations and caveats”. In: Applied Animal Behaviour Science 163, pp. 1–18.

Sarewitz, D. and R. Pielke (1999). “Prediction in science and policy”. In: Technology in Society 21.2, pp. 121–133.

Stan Development Team et al. (2018). CmdStan: the command-line interfact to Stan. Version *2.23.0.* url: http://mc-stan.org.

Stan Development Team (2018). Stan modeling language user’s guide and reference manual, version *2.18. 0*. url: http://mc-stan.org.

Taylor, K. D. and D. S. Mills (2006). “The development and assessment of temperament tests for adult companion dogs”. In: Journal of Veterinary Behavior 1.3, pp. 94–108.

Tseng, C.-h. et al. (2016). “Longitudinal data analysis with non-ignorable missing data”. In: Statistical Methods in Medical Research 25.1, pp. 205–220.

Valsecchi, P. et al. (2011). “Temperament test for re-homed dogs validated through direct behavioral observation in shelter and home environment”. In: Journal of Veterinary Behavior 6.3, pp. 161–177.

Van der Borg, J. A., W. J. Netto, and D. J. Planta (1991). “Behavioural testing of dogs in animal shelters to predict problem behaviour”. In: Applied Animal Behaviour Science 32.2–3, pp. 237–251.

Vonesh, E. F., T. Greene, and M. D. Schluchter (2006). “Shared parameter models for the joint analysis of longitudinal data and event times”. In: Statistics in Medicine 25.1, pp. 143–163.

Wang, C.-C. and W.-C. Lee (2019). “A simple method to estimate prediction intervals and predictive distributions: Summarizing meta-analyses beyond means and confidence intervals”. In: Research Synthesis Methods 10.2, pp. 255–266.

