## Supplementary Materials for "Predicting individual shelter dog behaviour after adoption using longitudinal behavioural assessment: a hierarchical Bayesian approach"

**2 Model description** **5**

**3 Variation across contexts** **8**

**4 Repeatability** **9**

**5 Random effect correlations** **10**

**6 Examples of individual-level posterior predictions** **10**

### 1 Behavioural codes

Below is a list of all behavioural codes (categorised by colour) used to record dog behaviour. See the post-rehoming telephone questionnaire for which codes are used to score behaviour in particular contexts and how they are ordered.

#### 1.1 Green codes

- **Friendly**: dog initiates interaction with people or dogs in an appropriate social manner. A dog in this category may display an appropriate level of reprimand towards other dogs but must be predominantly social.
- **Relaxed**: free from tension and anxiety, body movements should be fluid. The dog responds positively to his environment (interactive, free from stress or overstimulation).
- **Excitable**: the dog will have an enthusiastic attitude with animated interaction, showing behaviours such as jumping up, mouthing, an inability to stand still, playful towards people, dogs and resources.
- **Playful**: the dog has an interactive attitude towards wanting to engage in activities or social games.
- **Vocal**: dog barks, whines, howls or makes any other vocalisation in a non-aggressive manner. The dog is generally vocal, not in response to the presence or absence of people or other dogs.
- **Depressed**: the dog will generally be reserved and withdrawn, reluctant to interact with people/dogs or their environment.
- **Independent**: the dog doesn't actively seek interaction with other dogs/people however is relaxed in the present of people/other dogs.
- **Not motivated**: the dog has no interest in interacting with toys.
- **Not Eating Meal**: the dog is not eating any food that is given. Staff member will need to state what different types of food have been offered, how long he has had access to it and at what time of the day. N.B. some anxious dogs will only eat overnight whilst settling in.
- **Not Eating Treat**: the dog is not interested in eating any treats but is eating his main meal. Staff member will need to state what treats the dog has been offered and in which situation.
- **Submissive**: the dog is showing appeasing and or nervous behaviours with features such as low body posture rolling over and other calming signals.
- **Stressed**: the dog will be showing more than one of the 'stress behaviours' which may include panting, pacing, yawning, etc.
- **Unsure**: the dog can be apprehensive and reluctant to seek interaction with dogs/people or be concerned by the environment.

#### 1.2 Amber codes

- **Stressed** +: the dog is showing high frequency/intensity of the stress behaviours, and may also include dribbling, stereotypic behaviours, stress vocalisations, constant shedding, trembling, destructive behaviours etc.
- **Uncomfortable avoids**: the dog has a tense and stiff posture and/or shows anxious behaviours (potentially giving calming or distance increasing/decreasing signals) while trying to move away from the situation (another dog, person, handling, traffic etc.).
- **Submissive** +: the dog is showing high intensity of the submissive behaviours such as submissive urination, a reluctance to move, or is frequently overwhelmed by the environment or interactions.
- **Uncomfortable Static**: the dog has tense and stiff posture and/or shows anxious behaviour (potentially giving calming signals or distance increasing/decreasing signals) but doesn't attempt to move away from the situation (another dog, person, resources, handling, traffic etc.).
- **Uncomfortable Approaches:** the dog has tense and stiff posture and/or shows anxious behaviour (potentially giving calming signals or distance increasing/decreasing signals) and approaches the situation (e.g. another dog, person, traffic etc.)
- **Sexual**: the dog will be showing mounting behaviours or is especially interested in the genital area (e.g. sniffing licking) towards dogs but can be distracted away.
- **Sexual** +: the dogs will be showing high frequency/intensity of the sexual behaviours not offering any other social interaction, very difficult to distract.
- **Chases**: the dog has a tendency to be motivated and wants to run after the movement of people/dogs/joggers and will become stimulated by this.
- **Focused**: the dog is fixated on stimulus (must state what). It is difficult to get their attention back on to the handler. The dog may have lowered posture or stalk them but is not vocal.
- **Playful** +: the dog will be showing high frequency/intensity of the playful behaviours, don't seem to have any boundaries within their interaction (includes rude).
- **Reactive to People Non Aggressive**: barks, whines, howls and/or play growls when seeing/meeting other people, potentially pulling or lunging towards them (state radius).
- **Reactive to Dogs Non Aggressive**: barks, whines, howls and/or play growls when seeing/meeting other dogs, potentially pulling or lunging towards them (state radius).

- **Depressed** +: the dog is in total shut down and is not responsive to anything within its environment.

#### 1.3 Red codes

- **Stressed** ++: the dog is showing very high frequency/intensity of the stress behaviours and/or self-mutilation, injuring themselves in the environment. These behaviours impact significantly on the dogs' welfare.
- **Chases** +: the dog is showing high frequency/intensity of the chase behaviours very fixated on the movement and trying to grab or becoming reactive towards. It is likely that this behaviour is practiced and the dog can't be distracted from doing it.
- **Overstimulated**: the dog is showing high intensity of the excitable behaviours and/or grabbing, body barging, nipping etc.
- **Uncomfortable Static** +: the dog has given a freeze in response to a particular situation.
- **Reactive to People Aggressive**: growls, snarls, shows teeth and/or snaps when seeing/meeting other people, potentially pulling or lunging towards them (state radius).

### 2 Model description

As summarised in the main text, we specified a custom mixture model for the probability of different codes ($c = \text{missing}, \text{green}, \text{amber}, \text{red} = 0, 1, 2, 3$) for case $i$ (either the days after arriving at the shelter or the days after adoption), dog $j$ and context $k$:

$$p(y^c_{ijg}) = \begin{cases} \psi_{ijg} & \text{if } y_{ijg} = 0 \\ (1 - \psi_{ijg})\left[\kappa + (1-\kappa)\pi^c_{ijg}\right] & \text{if } y_{ijg} = 1 \\ (1 - \psi_{ijg})(1-\kappa)\pi^c_{ijg} & \text{if } 1 < y_{ijg} \leq 3 \end{cases} \quad (1)$$

This custom likelihood function is defined in the log probability mass function `ordinal_hurdle_one_inflated_lpmf` in the Stan model file (https://github.com/goold-newberry-lba/analysis-code/jhb-ordinal-hurdle-model.stan) as:

```
/* log-PDF for a single response from the custom ordinal-hurdle model with one-inflation
 *     y:    ordinal data point
 *     pi: linear predictor for the hurdle model
 *     theta: vector of ordinal category probabilities
 *     kappa: probability of one-inflation
*/
real ordinal_hurdle_one_inflated_lpmf(int y, real eta, vector pi, real kappa){

    real lp = 0;

    if(y == 0){
      lp += bernoulli_logit_lpmf(1 | eta);
    }

    if(y == 1){
      vector[2] lp1;
      lp1[1] = bernoulli_logit_lpmf(0 | eta) + log(kappa);
      lp1[2] = bernoulli_logit_lpmf(0 | eta) + log1m(kappa) + categorical_lpmf(y | pi);
      lp += log_sum_exp(lp1);
    }

    if(y > 1){
      lp += bernoulli_logit_lpmf(0 | eta) + log1m(kappa) + categorical_lpmf(y | pi);
    }

    return lp;
  }
```

The above code defines a log-probability accumulator variable `lp`, and adds the relevant probability components to the latter variable depending on the data point `y` passed to the function. When `y==1`, the mixture components are defined with probability $\kappa$, and we use the `log_sum_exp` function to safely take the log of the summed probabilites of the mixture components.

The vector `pi` passed to the above function holds the probabilities of green, amber and red codes, which are determined based on a cumulative ordinal probit model. For the shelter sub-model, this is defined as:

$$\pi^c_{ijg_s} = \Phi\Big(\frac{\theta_{c_s} - \mu_{ijg}}{\sigma}\Big) - \Phi\Big(\frac{\theta_{c-1_s} - \mu_{ijg}}{\sigma}\Big) \tag{2}$$

and, similarly, for the post adoption sub-model as:

$$\pi^c_{ijg_a} = \Phi\Big(\frac{\theta_{c_a} - \zeta_{ijg}}{\epsilon}\Big) - \Phi\Big(\frac{\theta_{c-1_a} - \zeta_{ijg}}{\epsilon}\Big) \tag{3}$$

where the $s$ and $a$ subscripts denote the shelter and post-adoption parameters, respectively. The $\mu$ and $\zeta$ parameters hold the latent behavioural scale means for the shelter and post adoption sub-models, which are specified as the following additive functions:

$$\mu_{ijk} = \alpha + r^1_j + r^1_g + r^1_{j\times g} + \big(\beta + r^2_j + r^2_g + r^2_{j\times g}\big)x_1 + \mathbf{X_1 B} \tag{4}$$

$$\zeta_{ijk} = \delta + r^4_j + r^4_g + r^4_{j\times g} + \big(\gamma + r^5_j + r^5_g + r^5_{j\times g}\big)x_2 + \mathbf{X_2 \Gamma} \tag{5}$$

where the random effects are denoted by $r$ (see below), $x_{1,2}$ denote the days after arrival and days after adoption with coefficients $\beta$ and $\gamma$, respectively, and $\mathbf{X_{1,2}}$ are the matrices of dog-level predictor variables with coefficient vectors $\mathbf{B}$ and $\mathbf{\Gamma}$, which are detailed in Table 1 and in section 'Predictor variables' in the statistical methods of the main text.

The probability of missing data in each sub-model is modelled as:

$$y^c_{ijg_s}\big|_{c=0} \sim \text{Bernoulli}(logit^{-1}(\eta_{ijg})) \tag{6}$$

$$y^c_{ijg_a}\big|_{c=0} \sim \text{Bernoulli}(logit^{-1}(\nu_{ijg})) \tag{7}$$

where $y_{ijg} = 0$ denotes a missing data point. The parameters $\eta$ and $\nu$ are additive functions on the linear log-odds scale, hence the use of the inverse logit link above to map to the probability space required by the Bernoulli likelihood. The additive functions were defined as:

$$\eta_{ijg} = \alpha_{miss} + r^3_j + r^3_g + r^3_{j\times g} + \beta_{miss}x_1 + \mathbf{X_1 B_{miss}} \tag{8}$$

$$\nu_{ijg} = \delta_{miss} + r^6_j + r^6_g + r^6_{j\times g} + \gamma_{miss}x_2 + \mathbf{X_2 \Gamma_{miss}} \tag{9}$$

A key component of our model is the correlated random effect parameters between shelter and post adoption time periods, which defines the 'joint' parameterisation of our model. The correlations enable information to flow between shelter and post adoption sub-models. The random effects on the intercept and slope (days after arrival to the shelter/after adoption) parameters above are present for dogs, contexts and dogs × context combinations, each of which are packaged into a 6 × 6 covariance matrix. To speed up the MCMC computations, we use the 'non-centered' parameterisation of the random-effect parameters and a Cholesky factorisation of their correlation matrices using the LKJ prior distribution in Stan.

As can be seen in the model file, we use weakly-informative prior distributions to aid computation and inference. Predictor coefficients were all given standard normal (i.e. $Normal(0, 1)$) prior distributions, all intercepts were given $Normal(0, 5)$ prior distributions, all standard deviations were given half standard normal priors (i.e. constrained to be positive), and LKJ priors were given to the Cholesky factors of the correlation matrices. The ordinal thresholds parameters $\theta$ have uniform priors.

### 3 Variation across contexts

Figure S1 demonstrates the variation across contexts for the latent behavioural scores (intercepts and slopes) and missing data, for an average dog, at the shelter. Figure S2 displays the same information for the post-adoption reports. For the shelter estimates, the intercepts in Figure S1a and the values in Figure S1b are evaluated at 32.1 days after arrival, and the slopes in S1a illustrate the amount of behavioural change for every 73 days (1 standard deviation of days after arrival) spent at the shelter. The intercepts S2a and the values in Figure S2b are evaluated at 30.1 days post adoption, and the slopes in S2a illustrate the amount of behavioural change for every 11.6 days after adoption. The missing probabilities in Figure S2b are low (approximately 1%) because they represent the average probability of missing data within contexts across dogs with either one and two surveys. The mean missingness decreases to essentially zero for dogs with two surveys, and increases to approximately 45% for dogs with only one completed survey.

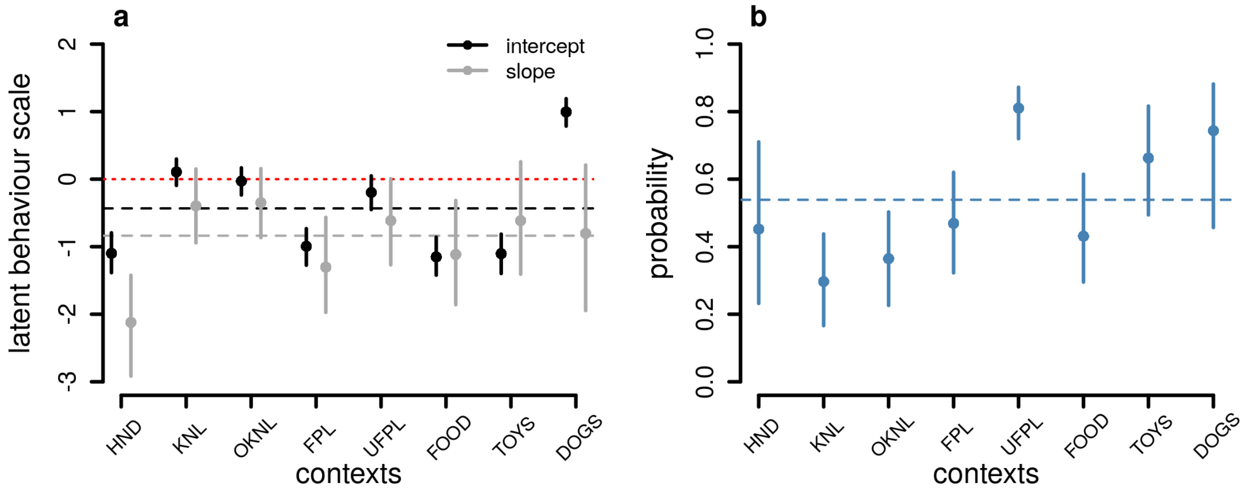

*Figure S1: Variation across contexts at the shelter (HND=handling, KNL=in kennel, OKNL=outside of kennel, FPL/UFLP=interactions with familiar/unfamiliar people, FOOD=eating food, TOYS/-DOGS=interactions with toys/dogs) for latent behavioural scores (panel a; higher intercept scores indicate higher chances of amber and red codes, more positive slopes indicate higher chances of amber and red codes through time) and the probability of missing data within surveys (panel b). The red dotted line in panel a highlights zero, while the dashed line in both panels indicates the average value across contexts.*

### 4 Repeatability

Repeatability is the proportion of behavioural variance attributable to between-individual differences. The model used here had four sources of behavioural variance (i.e. not including the missing data component) for both the shelter ($s$) and post adoption ($a$) sub-models: variation across dogs ($\sigma_j^{s,a}$), variation across contexts ($\sigma_g^{s,a}$), variation across dog $\times$ context combinations ($\sigma_{j\times g}^{s,a}$), and error variance ($\sigma$ or $\epsilon$, respectively). Therefore, we estimated repeatability across dogs, across contexts, and across dog $\times$ context combinations separately. For example, repeatability for dog $\times$ context combinations at the shelter was calculated as:

$$R_{j\times g}^{\text{shelter}} = \frac{(\sigma_{j\times g}^s)^2}{(\sigma_j^s)^2 + (\sigma_g^s)^2 + (\sigma_{j\times g}^s)^2 + \sigma^2} \tag{10}$$

and repeatability for contexts post adoption was calculated as:

$$R_g^{\text{adoption}} = \frac{(\sigma_g^a)^2}{(\sigma_j^a)^2 + (\sigma_g^a)^2 + (\sigma_{j\times g}^a)^2 + \epsilon^2} \tag{11}$$

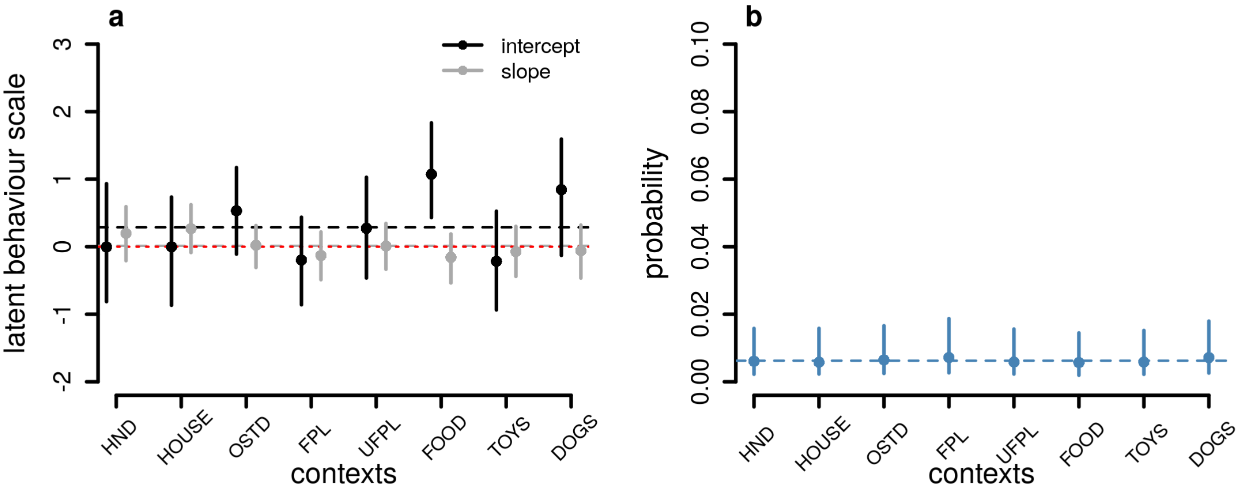

*Figure S2: Variation across contexts post-adoption (HND=handling, KNL=in kennel, OKNL=outside of house, FPL/UFLP=interactions with familiar/unfamiliar people, FOOD=eating food, TOYS/-DOGS=interactions with toys/dogs) for latent behavioural scores (panel a; higher intercept scores indicate higher chances of amber and red codes, more positive slopes indicate higher chances of amber and red codes through time) and the probability of missing data within contexts, within surveys (panel b). The red dotted line in panel a highlights zero, while the dashed line in both panels indicates the average value across contexts.*

### 5 Random effect correlations

The full list of random effect correlations estimated by the above model is presented in Table S1 found at https://github.com/cmgoold/goold-newberry-lba/tree/master/paper.

### 6 Examples of individual-level posterior predictions

A sample of 50 dogs' posterior predictions, each within a random context, is shown at https://github.com/cmgoold/goold-newberry-lba/tree/master/analysis-code/sample-dog-plots.
